# A novel transposable element in lampreys that is similar in sequence to *Tol2* of teleost fishes

**DOI:** 10.64898/2026.01.12.699174

**Authors:** Atsuo Iida, Manuel Ángel Pombal, Akihiko Koga, Eiichi Hondo

## Abstract

*Tol2* is one of the few DNA-based transposable elements that reside in vertebrate genomes and retain transposition activity. It was first identified in the medaka fish (*Oryzias latipes*) and subsequently in another species of the genus *Oryzias*, which belongs to the order Beloniformes. Transposable elements similar in sequence to *Tol2* also occur in several species of the order Cypriniformes, including the common carp. The low sequence variation of these elements relative to that of a nuclear gene provided evidence for past horizontal transfer events among host organisms. We collectively refer to *Tol2* and these related elements as the *Tol2* family. Here, we identified a novel transposable element in two European lamprey species (*Lampetra fluviatilis* and *L. planeri*), which we named *Tel2*. Lampreys belong to the jawless vertebrate lineage (infraphylum Agnatha), whereas all previously known host species of the *Tol2* family are jawed vertebrates (infraphylum Gnathostomata). The nucleotide sequence similarity between *Tol2* and *Tel2* was higher than that observed for the nuclear gene, indicating at least one horizontal transfer event between the two infraphyla. Thus, the *Tol2* family now includes *Tel2* as a new member. These results indicate that *Tol2* family elements have undergone multiple horizontal transfer events across a wide range of host organisms, including both jawless and jawed vertebrates. Because some members of this family still retain transposition activity, they provide an excellent and unique model for studying the role of horizontal transfer in the long-term persistence of DNA-based transposable elements.

## Introduction

DNA-based transposable elements are repetitive sequences that change their chromosomal locations in a cut-and-paste manner through the activity of the transposase enzyme (Wells 2020). It is generally thought that DNA-based transposable elements lose their transposition activity over time in the organisms that host them (Hartl 1997; Miskey 2005; Huang 2012). The reason is that active elements act as mutators in the host genome. Although transposition may result in beneficial mutations for the host, deleterious mutations are considered far more frequent. This leads to reduced fitness in host individuals or gametes carrying active element copies compared with those lacking such copies. In addition to this negative selection imposed by the elements themselves, host organisms may evolve proteins or mechanisms that suppress transposition activity, as has been observed for the *Drosophila* P and *hobo* elements (Bingham 1982; McGinnis 1983). Horizontal transfer is considered a key factor in the long-term survival of DNA-based transposable elements because selfish amplification may be permitted in a new host, at least for a short period after invasion (Lohe 1995; Schaack 2010). From this perspective, *Tol2* (transposable element of *<u>O</u>ryzias latipes*, medaka fish, no. <u>2</u>) is an intriguing DNA-based transposable element.

*Tol2* was identified 30 years ago in medaka as the first active DNA-based transposable element in a vertebrate (Koga 1996). It belongs to the *hA*T (*<u>h</u>obo*/*<u>A</u>ctivator*/<u>T</u>am*3*) superfamily (Atkinson 1993). This element is present on medaka chromosomes in 10–20 copies per haploid genome (Koga 1999a). Each copy is approximately 4.7 kb in length and contains 17- and 19-bp terminal inverted repeats (TIRs). The element is flanked by 8-bp target site duplications (TSDs), which are tandem repeats derived from the host chromosomal sequence at the insertion site. These two structural features (TIRs and TSDs) are common to many DNA-based transposable elements. *Tol2* also carries another set of repeat sequences unique to this element: 302- and 303-bp internal inverted repeats (IIRs). The *Tol2* element contains four exons that encode its transposase. Several mutations in host genes have been shown to result from the insertion or excision of *Tol2* (Iida 2004, 2005; Koga 2006).

In the context of horizontal transfer, *Tol2* stands out for its copy composition: its copies are highly homogeneous in sequence, and virtually all are autonomous. To our knowledge, no other DNA-based transposable element has been reported to exhibit a level of sequence homogeneity comparable to that of *Tol2*. This feature led to the inference that the element was introduced into the medaka genome relatively recently (Koga 1999a). Thus, *Tol2* provides a unique model for investigating three key issues: (1) how the element will decay over time, (2) the possibility of horizontal transfer to another organism, and (3) the history of past horizontal transfer events. Of these, issues (1) and (2) concern future events and are therefore extremely difficult to examine unless they occur at a rapid rate, as observed for the *Drosophila* P element. The present study addresses issue (3), focusing on the taxonomic range of host organisms and the number of horizontal transfer events that occurred before *Tol2* reached the medaka genome.

In addition to medaka, *Tol2* has been identified in the genome of the Hainan medaka (*O. curvinotus*) (Koga 2000). Nucleotide and amino acid sequences were compared with those of the *tyrosinase* (*TYR*) gene, a nuclear gene that is relatively highly conserved because of its essential role in host survival (Sato 2001). Sequence similarity was higher for *Tol2* than for *TYR*, leading to the conclusion that horizontal transfer occurred either between the two *Oryzias* species or independently into both species from a common source. The genus *Oryzias* belongs to the order Beloniformes. Elements similar in sequence to *Tol2* are also present in numerous species of the order Cypriniformes, including the common carp (*Cyprinus carpio*) and the emerald dwarf rasbora (*Danio erythromicron*) (Jiang 2012; Ishiyama 2017). Based on sequence analyses, these elements were considered to share a common origin with *Tol2*. In the present paper, we collectively refer to *Tol2* and these related elements as the *Tol2* family. Comparisons of sequence similarity between *Tol2* family elements and the *TYR* gene consistently showed higher similarity among the *Tol2* family elements, indicating that horizontal transfer of these elements occurred between the two orders. The occurrence of multiple horizontal transfer events led us to expect that additional organisms might harbor members of the *Tol2* family.

For this purpose, we searched genome sequence databases. However, we obtained no positive results for years. During the early stages of genome database development, contig sequences were assembled from short sequencing reads. Repetitive sequences, particularly highly homogeneous ones, were often underrepresented in databases generated using this approach (Koga 2012). This technical limitation hindered our ability to identify *Tol2* family elements through database surveys. More recently, advanced long-read sequencing technologies have been developed (Jain 2018; Miga 2020), including nanopore sequencing and single-molecule real-time sequencing, enabling the rapid generation of high-quality whole-genome sequences. Telomere-to-telomere and chromosome-level genome assemblies are now available for many species (Logsdon 2020; Li 2024). The application of these technologies has expanded to a wide range of organisms, including nontraditional model animals. In this study, we surveyed the latest genome databases and identified a novel *Tol2* family element in the lamprey genome.

## Results

### 1. Initial survey

We performed BLAST searches using the entire *Tol2* sequence as the query against all sequences deposited in the NCBI database. Among the top 100 hits, 95 were derived from animals and five were artificial sequences (Figure 1, Table S1). The animal-derived sequences comprised 51 from medaka, the original host species of *Tol2*; 23 from cyprinid fishes, another host taxonomic group previously reported to harbor *Tol2*; five from *Amia ocellicauda*, a freshwater fish; and 16 from lampreys. Whereas the fish species, excluding lampreys, are jawed vertebrates (infraphylum Gnathostomata), lampreys belong to the jawless vertebrates (infraphylum Agnatha). To the best of our knowledge, no transposable element similar in sequence to *Tol2* has previously been reported in lampreys. Given the distant evolutionary relationship between jawed and jawless vertebrates, we conducted further analyses of the lamprey-derived sequences.

**Figure 1.**
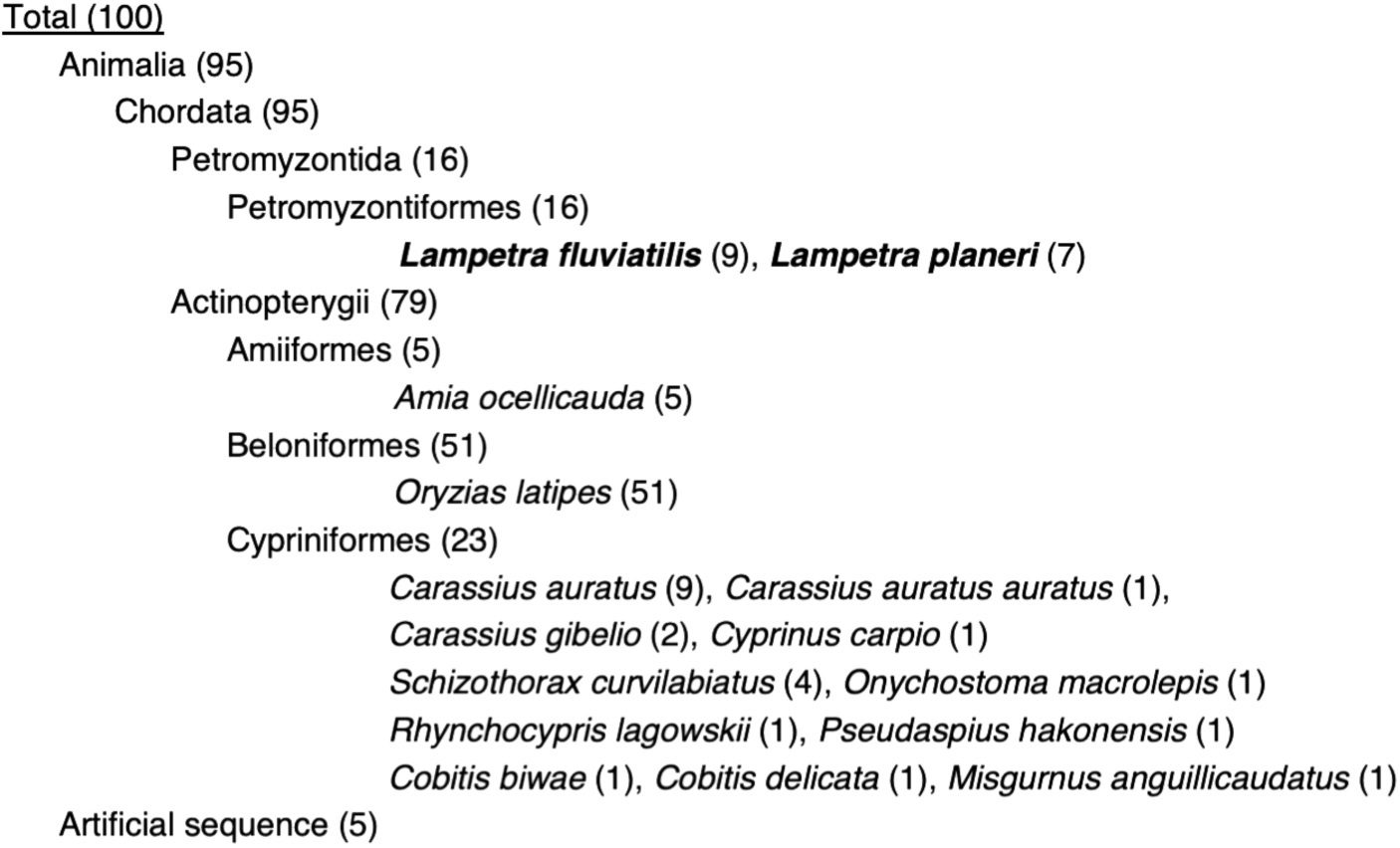
BLAST survey for the identification of *Tol2*-like sequences. The illustration shows the taxonomic classification of the species identified in the initial BLAST survey using the full-length *Tol2* sequence as the query. The numbers in parentheses indicate the total number of subject sequences identified in each taxonomic group or species. The complete list of subject sequences is provided in Table S1.

### 2. Secondary survey

Restricting the search to lampreys (Taxonomy ID: 7745), we conducted another round of surveys using 150-bp nucleotide blocks from the left or right end of the *Tol2* sequence as queries. Searches with the left-terminal query detected 18 sequences in *L. fluviatilis* and 17 in *L. planeri*. Searches with the right-terminal query identified 16 sequences in *L. fluviatilis* and 13 in *L. planeri* (Figure 2). No significant hits were detected in lamprey species other than *L. fluviatilis* and *L. planeri*. Subsequent chromosomal mapping indicated that the *L. fluviatilis* genome assembly contains at least six *Tol2*-like sequences with both TIRs, nine with the left TIR only, and six with the right TIR only (Table S2). Using the same criteria, seven sequences with both TIRs, ten with the left TIR only, and six with the right TIR only were identified in the *L. planeri* genome assembly (Table S3). Thus, the *L. fluviatilis* individual used as the DNA source contained at least 21 *Tol2*-like sequences per haploid genome, whereas the *L. planeri* individual contained at least 23 such sequences per haploid genome. We assigned an identifier (ID) to each sequence (Tables S2 and S3).

**Figure 2.**
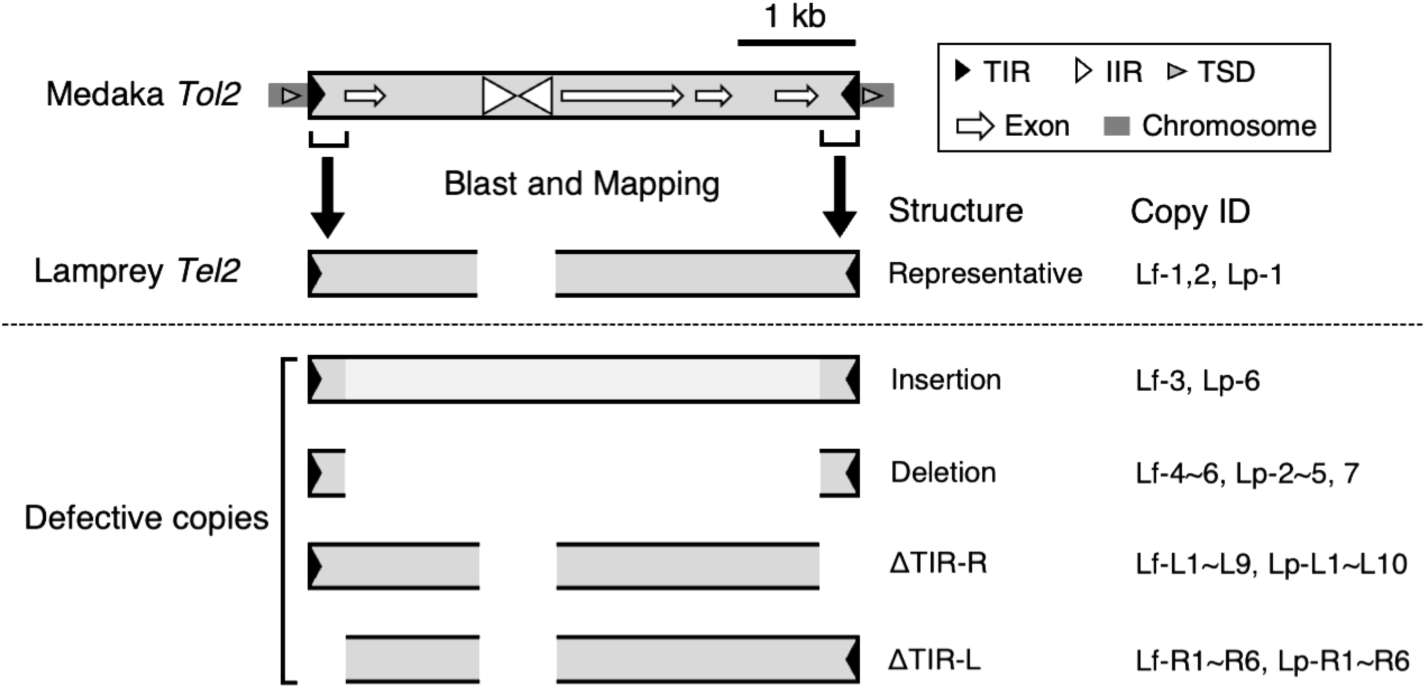
Assembly of lamprey *Tel2* sequences. *Tel2* terminal sequences were identified through homology searches using the terminal sequences of *Tol2* as queries. The corresponding internal *Tel2* sequences and adjacent chromosomal regions were then associated with these terminal fragments, enabling reconstruction of the full-length sequence of each copy. In total, 44 *Tel2* sequences were identified, mapped to their chromosomal loci, and assigned unique identifiers (IDs) (Tables S2 and S3). TIR, terminal inverted repeats; IIR, internal inverted repeat; TSD, target site duplication; Lf, *L. fluviatilis*; Lp, *L. planeri*.

All mapped sequences, except for Lp-L6, were also present at the corresponding chromosomal positions in the alternative haplotype assembly. We assumed that these lamprey sequences are DNA-based transposable elements sharing a common origin with *Tol2*. Based on this assumption, the Lf-1 copy, which is 4,032 bp in length, most closely resembles the sequence and structure of the ancestral element because it exhibits relatively high sequence similarity to *Tol2* across its entire length. Lf-2 from *L. fluviatilis* and Lp-1 from *L. planeri* were both 3,981 bp long and contained TIRs together with an internal sequence similar to that of Lf-1. Therefore, these copies were regarded as the next most ancestral sequences. The remaining sequences lacked one or more *Tol2*-specific motifs and/or were substantially shorter or longer than Lf-1. Based on the assumption that these elements share a common origin with *Tol2*, we named the *Tol2*-like sequences identified in lampreys *Tel2* (transposable element in <u>E</u>uropean lampreys, *Tol<u>2</u>*-related).

### 3. Terminal inverted repeats

The presence of terminal inverted repeats (TIRs) is one of the defining structural features of DNA-based transposable elements. TIRs are thought to serve as substrates, or components of substrates, for the transposase enzyme (Kidwell 2005; Tafalla 2006). Their sequences are typically unique to a particular transposable element family or to closely related families. *Tol2* possesses a 17-bp left TIR and a 19-bp right TIR. With this in mind, we first analyzed the terminal regions of *Tel2* for the presence of TIRs and then compared them with those of *Tol2* (Figure 3A). In the *L. fluviatilis* genome, six *Tel2* copies spanning from the left to the right end had been identified. Similarly, seven such copies were identified in *L. planeri*. All 13 *Tel2* copies were found to contain 17-bp left and 19-bp right TIRs (Figure 3B). Their TIR sequences were identical to those of *Tol2*, except for a single-base mismatch in the Lf-2 and Lp-1 copies. We also analyzed *Tel2* copies containing only one terminal region. Their predicted TIR sequences were identical or nearly identical to those of *Tol2* (Figure S1).

**Figure 3.**
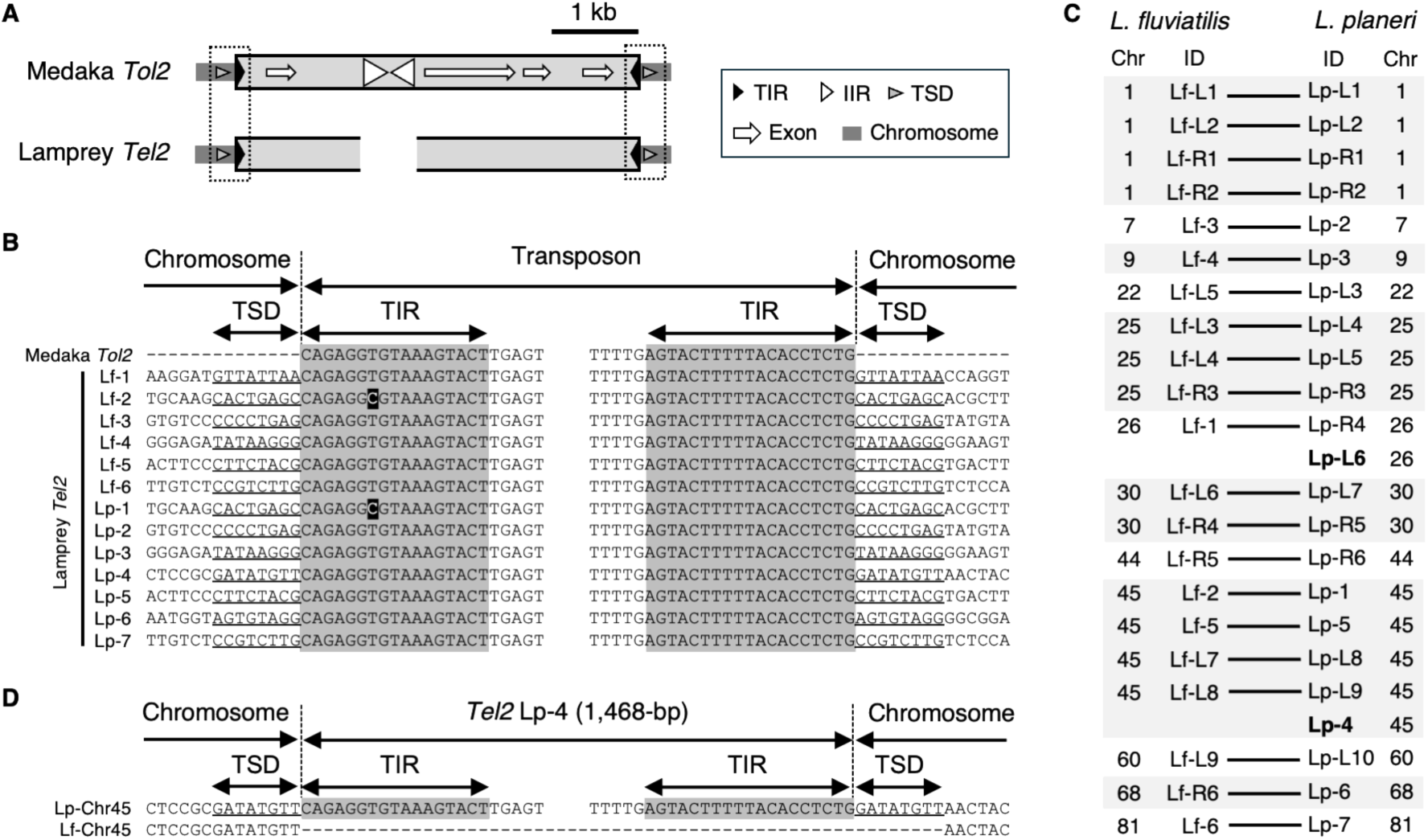
*Tel2* terminal structures and adjacent host genomic sequences. **A**. Overview of the *Tel2* termini and the adjacent host genomic sequences. **B**. Comparison of the terminal sequences of the *Tol2* element and the 13 *Tel2* copies containing both TIRs. Shaded bases indicate nucleotide substitutions relative to *Tol2*. Information on copies containing only one terminal region is provided in Figure S1. **C**. Correspondence of *Tel2* insertion loci between the chromosomes of *L. fluviatilis* and *L. planeri*. No corresponding copies were identified in *L. planeri* for the two IDs shown in bold. **D**. Comparison of the Lp-4 insertion site on chromosome 45 of *L. planeri* with the corresponding locus on chromosome 45 of *L. fluviatilis*, which lacks the *Tel2* insertion. TIR, terminal inverted repeat; IIR, internal inverted repeat; TSD, target site duplication; Lf, *L. fluviatilis*; Lp, *L. planeri*.

### 4. Target side duplication

Many transposable elements are known to generate target site duplications (TSDs) upon insertion into chromosomes (Kidwell 2005). In the case of *Tol2*, 8-bp TSDs are generated (Koga 1996). All 13 *Tel2* copies that contained both terminal regions were found to be flanked by identical 8-bp sequences, which were considered to represent TSDs (Figure 3B). This analysis could not be applied to *Tel2* copies containing only one terminal region (Figure S1).

### 5. Chromosomal location of *Tel2* copies

By analyzing longer flanking sequences, we mapped the *Tel2* copies, including those containing both terminal repeats as well as those with only one terminal region, onto the lamprey chromosomes (Figure 3B; Tables S2 and S3). Their chromosomal locations corresponded between *L. fluviatilis* and *L. planeri*, with the exception of two copies found only in the latter species (Figure 3C). Thus, most *Tel2* copies were shared between the two species, although the nucleotide sequences and lengths of the corresponding copies were not always identical. The only exceptions to this shared distribution were Lp-4 and Lp-L6. No counterpart of Lp-L6 was identified in the *L. fluviatilis* genome. Likewise, no counterpart of Lp-4 was found at the corresponding position on chromosome 45 of *L. fluviatilis* (Figure 3D). Instead, a single “GATATGTT” sequence was present at that position, identical to the 8-bp sequence duplicated on either side of the Lp-4 copy.

### 6. Absence of internal repeats

Comparison of Lf-1 with *Tol2* revealed a notable structural difference: an internal region of approximately 0.7 kb present in *Tol2* was absent from Lf-1 (Figure 4A). This feature was shared by all *Tel2* copies. The boundaries defining the beginning and end of the missing region were identical among the *Tel2* copies (Figure 4B). In the *Tol2* sequence, this region corresponds to the internal inverted repeats (IIRs), a structural feature unique to *Tol2* among members of the *hA*T superfamily.

**Figure 4.**
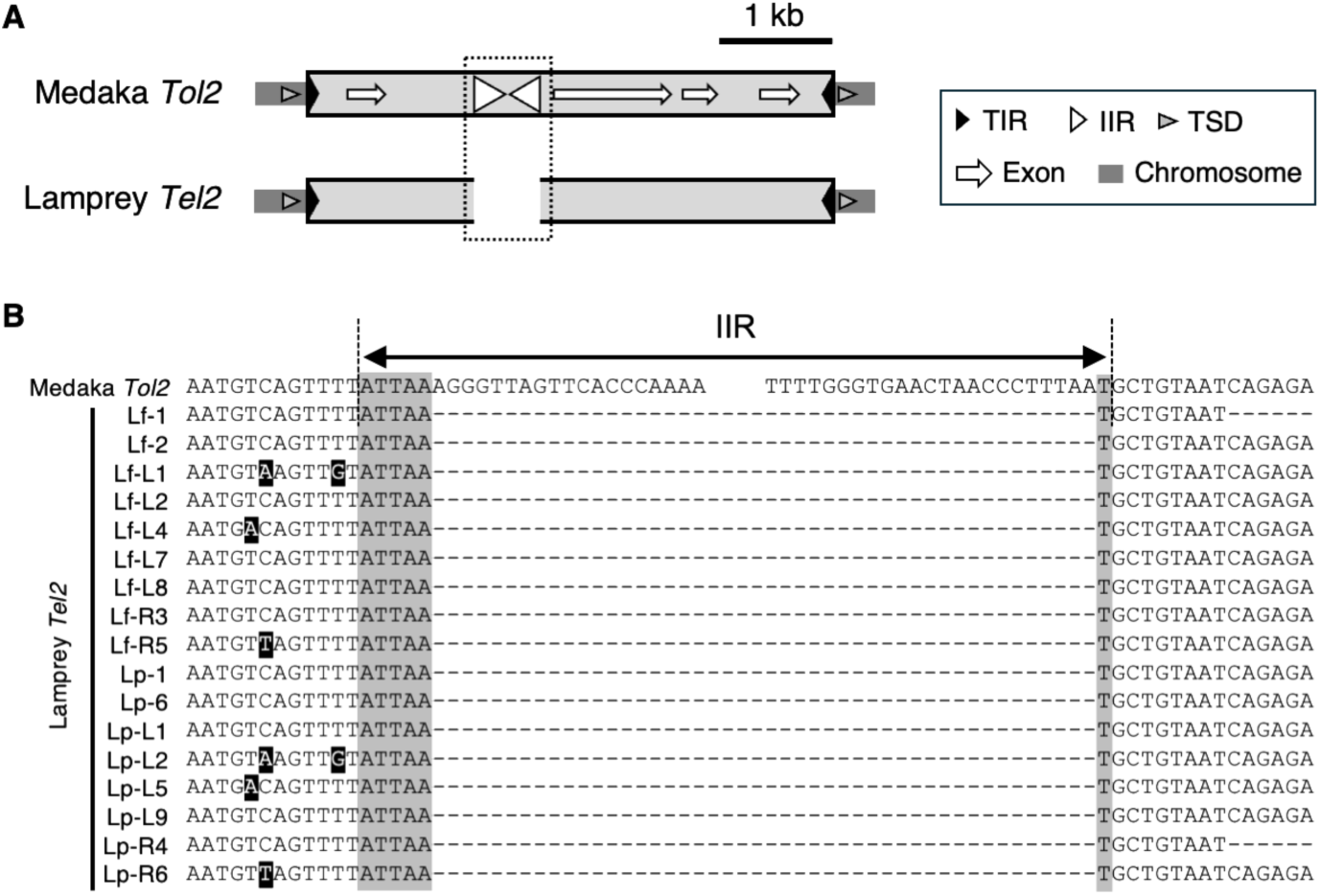
Absence of the IIRs in the *Tel2* internal region. **A**. Overview of the *Tel2* internal region corresponding to the location of the IIRs in the *Tol2* element. **B**. Comparison of the internal sequences of the *Tol2* element and 17 *Tel2* copies. An ATTAAT sequence is present in the *Tel2* internal region at the position corresponding to the IIRs in the *Tol2* element. TIRs, terminal inverted repeats; IIR, internal inverted repeat; TSD, target site duplication; Lf, *L. fluviatilis*; Lp, *L. planeri*.

### 7. Open reading frame

Autonomous DNA-based transposable elements contain a gene encoding transposase, the enzyme that catalyzes the transposition of both autonomous and nonautonomous copies. To identify coding sequences for transposase or other proteins that might be present in *Tel2*, we searched for open reading frames (ORFs) in *Tel2* copies containing TIRs at both ends. In *Tol2*, the transposase gene consists of four exons (Koga 1999b). Taking this into account, we defined an ORF as a reading frame beginning with any sense codon rather than being restricted to the canonical initiation codon (ATG). Figure 5 shows the distribution of ORFs of 300 bp or longer in all three reading frames on both DNA strands of the 13 *Tel2* copies examined (Lf-1 to Lf-6 from *L. fluviatilis* and Lp-1 to Lp-7 from *L. planeri*). Relatively short ORFs (300–600 bp) were observed in all 13 copies. ORFs longer than 600 bp were identified as follows: 909 bp in Lf-1; 1,128 bp in Lf-2; 1,944, 948, 690, and 2,031 bp in Lf-3; and 1,128 bp in Lp-1 (Tables S4 and S5).

**Figure 5.**
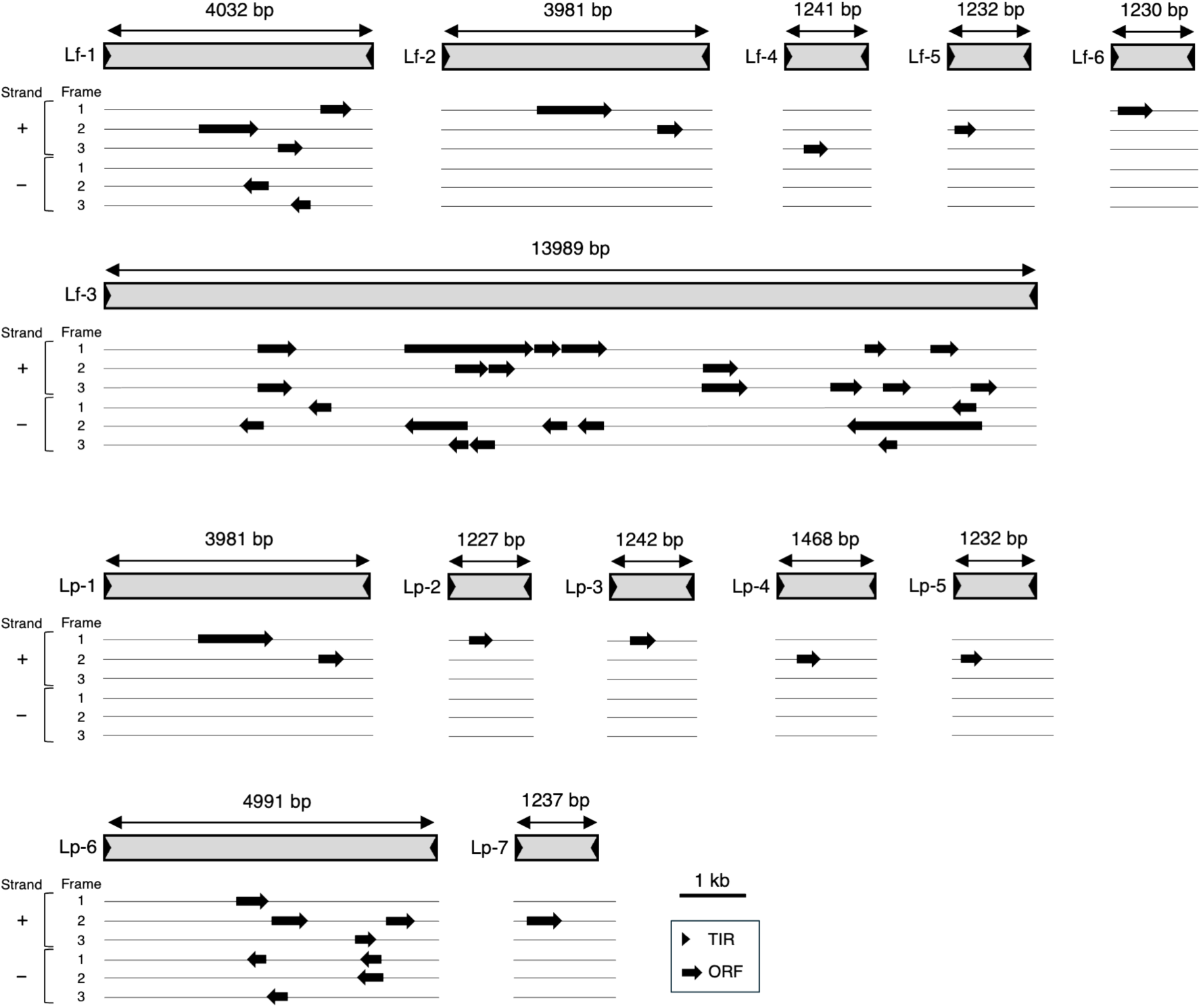
Open reading frames in *Tel2* copies. The illustration shows the open reading frames (ORFs) identified in the 13 *Tel2* copies containing both TIRs in *L. fluviatilis* and *L. planeri*. ORFs were defined as stop-to-stop coding sequences longer than 300 bp in all three reading frames on both DNA strands. TIR, terminal inverted repeat; Lf, *L. fluviatilis*; Lp, *L. planeri*.

### 8. Polymerase chain reaction (PCR)

We performed PCR analysis to verify the presence of *Tel2* sequences in the lamprey genome. The oligonucleotide primers were designed based on sequences shared between the *Tol2* element and the Lf-1 copy of *Tel2* (Figure 6A and Table S7). Genomic DNA was obtained from four *L. fluviatilis* individuals. Using a primer pair designed to anneal to the terminal regions of the transposons (P1a–P4b), we obtained amplicons of approximately 4.0 kb and 1.2 kb from the lamprey genome (Figure 6B). A 4.7-kb amplicon was obtained from the medaka sample. We next amplified specific regions using primer pairs targeting the left terminal region (P1a–P1b), the region adjacent to the internal inverted repeats (IIRs) (P2a–P2b), exon 2 of the transposase gene (P3a–P3b), and the right terminal region (P4a–P4b) of *Tol2*. All primer pairs except P2a–P2b amplified fragments of similar size in both species. The P2a–P2b primer pair amplified a 0.3-kb fragment from the lamprey specimens, which was 0.7 kb shorter than the corresponding fragment from medaka (Figure 6C). This finding is consistent with the genome database results, indicating that *L. fluviatilis* carries *Tel2* elements lacking the IIRs region present in the medaka *Tol2* element.

**Figure 6.**
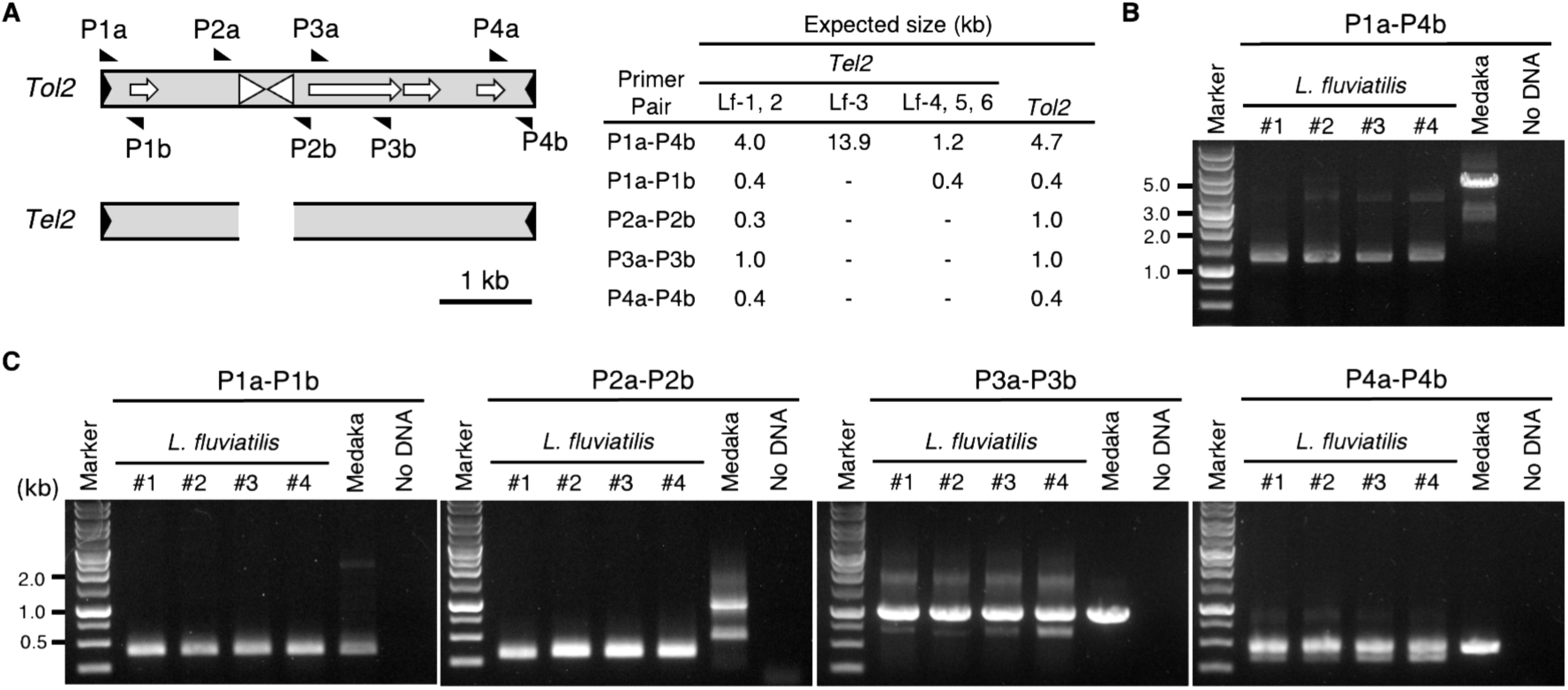
PCR analysis for the detection of *Tel2* sequences in lampreys. **A**. Positions of the primers and the expected amplicon sizes used in this study. The P1a and P4b primers were designed to amplify the full-length *Tol2* element and the *Tel2* copies (Lf-1, Lf-2, Lf-3, Lf-4, Lf-5, and Lf-6). Primer pairs used to amplify partial regions of the elements were designed based on the *Tol2* element and the Lf-1 and Lf-2 *Tel2* copies. The primer sequences are listed in Table S7. **B**. Agarose gel electrophoresis of PCR products amplified with the P1a–P4b primer pair, which targets the full-length elements. Four *L. fluviatilis* individuals (#1–#4) and one medaka were analyzed. The unexpected band is presumed to result from hairpin-loop formation mediated by the internal inverted repeats (IIRs) of the *Tol2* element. **C**. Agarose gel electrophoresis of PCR products amplified with the P1a–P1b, P2a–P2b, P3a–P3b, and P4a–P4b primer pairs, which target partial regions of the elements. The unexpected band observed with the P2a–P2b primer pair is presumed to result from hairpin-loop formation mediated by the internal inverted repeats (IIRs) of the *Tol2* element.

## Discussion

### 1. *Tel2* as a *Tol2* family transposable element

Through extensive genome database searches, we identified dispersed repetitive sequences in lampreys, which we named *Tel2*. We further confirmed the presence of *Tel2* by PCR analysis of lamprey genomic DNA. The *Tel2* copies identified in the genome databases—six from *L. fluviatilis* and seven from *L. planeri*—displayed the characteristic structural features of *hA*T family DNA-based transposable elements, including TIRs at both termini and 8-bp target site duplications (TSDs). The complete sequences of these *Tel2* copies, including their putative TIRs, exhibited high sequence similarity to the *Tol2* element from medaka. The Lp-4 copy provides particularly strong evidence supporting this conclusion. The Lp-4 sequence is flanked by two identical 8-bp nucleotide sequences (GATATGTT) on chromosome 45 of *L. planeri*. In contrast, the corresponding chromosomal position on chromosome 45 of *L. fluviatilis* lacks the *Tel2* insertion and contains only a single copy of this 8-bp sequence. This difference indicates that Lp-4 was inserted into the *L. planeri* genome by transposition, resulting in the formation of the TSD. Based on these findings, we conclude that *Tel2* is a member of the *Tol2* family, a group of transposable elements that share a common evolutionary origin with *Tol2*. Previously reported host organisms of *Tol2* family elements were limited to jawed vertebrates (infraphylum Gnathostomata) (Koga 1996; Jiang 2012; Ishiyama 2017). This is the first report of *Tol2* family elements in jawless vertebrates (infraphylum Agnatha), expanding the known host range of the *Tol2* family to encompass both major vertebrate lineages.

### 2. Horizontal transfer between jawless and jawed vertebrates

Jawless and jawed vertebrates are thought to have diverged during the Ordovician period or earlier (Escriva 2002; Wittbrodt 2002). To determine whether the sequence differences between *Tol2* and *Tel2* are consistent with this long evolutionary timespan, we compared them with the corresponding differences in the TYR gene. First, the ORF overlapping exon 2 of *Tol2* (1,131 bp) was compared with the corresponding *Tel2* ORFs: 909 bp in Lf-1 and 1,128 bp in Lf-2/Lp-1. Sequence alignment revealed high similarity between the *Tol2* and *Tel2* copies (Lf-1 or Lf-2/Lp-1), with nucleotide identities of 93% and 98% and deduced amino acid identities of 95% and 91%, respectively (Table 1). Next, we performed the same analysis using the complete coding sequences of *TYR* (1,623 bp in medaka, 1,608 bp in common carp, and 1,716 bp in both *L. fluviatilis* and *L. planeri*). The alignment scores for *TYR* were substantially lower than those for *Tol2* and *Tel2*, showing 47% nucleotide identity and 48% and 47% amino acid identity (Table 2). Thus, *Tol2* and *Tel2* exhibit a much higher degree of sequence similarity between these distantly related host groups than does *TYR*. We also compared these sequences with *Tol2*-cyp, a *Tol2* family element previously identified in cyprinids. A 4,714-bp *Tol2*-cyp sequence was isolated from the genome of *C. carpio* (common carp) based on a partial sequence reported previously (LC159588.1; Ishiyama 2017). A 1,140-bp region overlapping exon 2 of *Tol2* was then used as the query sequence for comparison. In addition, a 1,607-bp *TYR* coding sequence was retrieved from the *C. carpio* genome database and included in the analysis (Table S6). As in the lamprey comparison, sequence similarity was consistently higher among the transposase genes than among the *TYR* genes (Tables 1 and 2). Pairwise *dN*/*dS* ratios for *TYR* were all less than 1 (0.005–0.690, Table 3), as expected under purifying selection (Nei 1986; Li 1993). These results indicate that the *Tol2* family elements (medaka *Tol2*, lamprey *Tel2*, and carp *Tol2*-cyp) invaded their respective host lineages after those hosts had diverged (Figure 7A). This is the first demonstration of horizontal transfer of a member of the *hA*T superfamily between jawless and jawed vertebrates.

**Figure 7.**
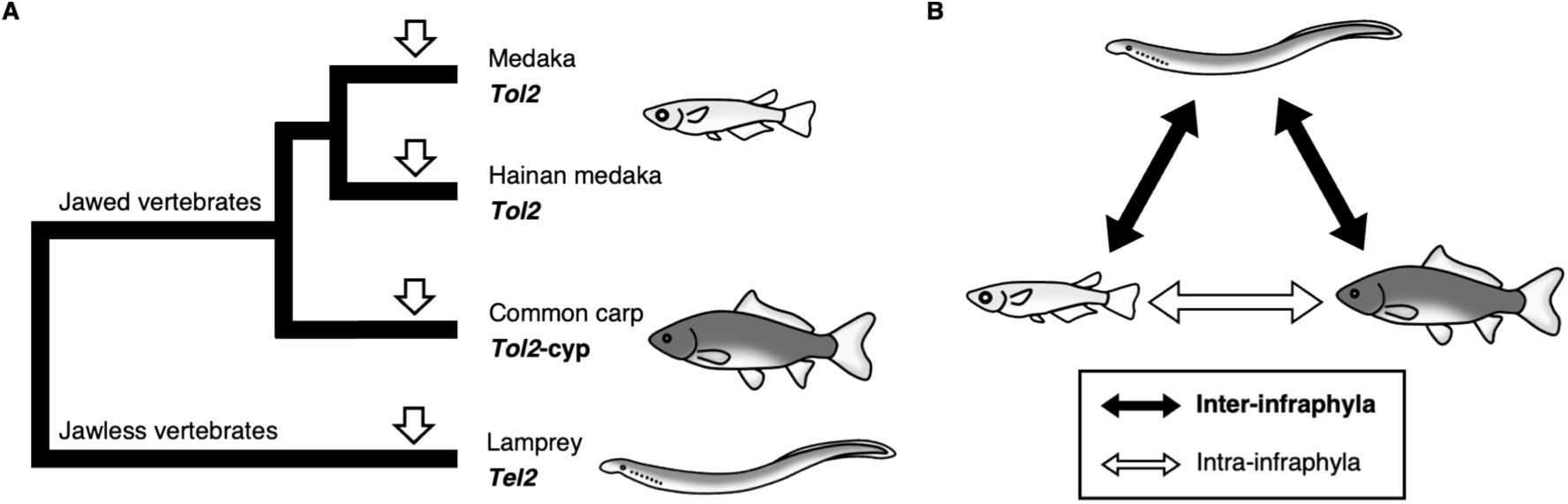
Horizontal transfer of *Tol2* family elements. **A**. Phylogenetic relationships among host organisms carrying *Tol2* family elements. *Tol2* family elements have been identified in both jawed and jawless vertebrate lineages. Arrows indicate the inferred invasion events of *Tol2* family elements. **B**. A hypothesis for the horizontal transfer of *Tol2* family elements among host organisms. *Tol2* family elements have been identified in three vertebrate lineages: lampreys, medakas, and carps. If the current distribution resulted from multiple horizontal transfer events, at least two independent transfer events would be required, including at least one transfer between the two vertebrate infraphyla.

**Table 1.** Sequence alignment scores (%) for selected ORFs calculated using CLUSTALW. Upper right: nucleotide sequences; lower left: deduced amino acid sequences.

|  | *Tol2* | *Tol2-cyp* | Lf-1 | Lf-2 / Lp-1 |
| --- | --- | --- | --- | --- |
| *Tol2* | – | 91 | 93 | 98 |
| *Tol2-cyp* | 87 | – | 88 | 91 |
| Lf-1 | 95 | 82 | – | 94 |
| Lf-2 / Lp-1 | 91 | 86 | 91 | – |

**Table 2.** Sequence alignment scores (%) for *TYR* coding sequences calculated using CLUSTALW. Upper right: nucleotide sequences; lower left: deduced amino acid sequences.

|  | Medaka | *C. carpio* | *L. fluviatilis* | *L. planeri* |
| --- | --- | --- | --- | --- |
| Medaka | – | 64 | 47 | 47 |
| *C. carpio* | 73 | – | 52 | 51 |
| *L. fluviatilis* | 48 | 50 | – | 99 |
| *L. planeri* | 47 | 50 | 99 | – |

**Table 3.** *dN*/*dS* ratios for *TYR* coding sequences calculated using MEGA X.

|  | *C. carpio* | *L. fluviatilis* | *L. planeri* |
| --- | --- | --- | --- |
| Medaka | 0.437 | 0.690 | 0.688 |
| *C. carpio* | – | 0.605 | 0.610 |
| *L. fluviatilis* | – | – | 0.005 |

### 3. Origin of internal inverted repeats

*Tel2* exhibits sequence similarity to *Tol2* throughout the region extending from the left TIR to the right TIR. However, one notable difference is found in an internal region: *Tol2* contains the internal inverted repeats (IIRs) sequence, whereas this region is absent from *Tel2*. A previous study (Izsvák 1999) reported that the *Tol2* IIRs originated through the insertion of a miniature inverted-repeat transposable element (MITE) named *Angel*. In that study, the consensus sequence of *Angel* was determined by comparing the nucleotide sequences of numerous *Angel* copies identified in genome sequence databases. The transposition mechanism of this MITE remains unclear. In rice, *mPing* was identified as a MITE, and its transposition was shown to be catalyzed by a transposase encoded by *Pong*, a longer autonomous element from which *mPing* is derived (Jiang 2003). As with most DNA-based transposable elements, *mPing* transposition generates target site duplications (TSDs) derived from the host chromosomal sequence. This example suggests that *Angel* transposition may likewise produce TSDs. If so, the currently defined *Angel* consensus sequence may include TSD sequences at its termini. In the present study, we obtained new evidence that helps define the boundary between the *Angel* element and the TSDs derived from the host chromosome. The *Tol2* IIRs is flanked by TTAA tetranucleotide sequences at both ends. In contrast, only a single TTAA sequence is present at the corresponding position in *Tel2*. This observation suggests that the terminal TTAA sequences included in the *Angel* consensus sequence are, in fact, TSDs generated during *Angel* transposition. As in the case of rice *mPing*, *Angel* may be mobilized by a yet-to-be-identified autonomous *Angel* element. The piggyBac transposable element is known to generate TTAA TSDs (Fraser 1996; Wu 2006). However, because terminal inverted repeat sequences generally serve as essential substrates for transposases, and because the terminal sequences of *Angel* and piggyBac are dissimilar, it is unlikely that *Angel* is mobilized by the piggyBac transposase. At present, the genetic mechanism by which the *Angel* element became incorporated into *Tol2* as the IIR remains unknown.

### 4. Occurrence of multiple horizontal transfer events

Horizontal transfer was first reported nearly 100 years ago in bacteria and has since been demonstrated in a wide range of multicellular organisms, including vertebrates (Griffith 1928; Crisp 2015). Unlike some RNA-mediated transposable elements, which share a common origin with retroviruses, DNA-based transposable elements lack the components required for intercellular transmission. Nevertheless, horizontal transfer has been documented for many DNA-based transposable elements (Schaack 2010; Zhang 2020). In the present study, we provide evidence for past horizontal transfer of *Tol2* family elements, including *Tel2*, across a broad range of vertebrates spanning the evolutionary boundary between jawless and jawed vertebrates. Below, we discuss the minimum number of horizontal transfer events required to explain the current distribution of *Tol2* family elements in vertebrate phylogeny.

Previous studies together with the present work show that *Tol2* family elements are distributed among three distantly related vertebrate lineages—lampreys, carps, and medakas—through horizontal transfer. The minimum number of horizontal transfer events required to explain this distribution is two (Figure 7B). In addition, one more event should be included based on a previous study: horizontal transfer between medaka and Hainan medaka, or independent transfer into both species from an unknown common source (Koga 2000). Furthermore, two additional events have been reported: one involving horizontal transfer within the order Cypriniformes (Ishiyama 2017), and another involving *Tgf2*, a *Tol2* family element that entered the goldfish (*Carassius auratus*) genome from an unknown source (Jiang 2012). Thus, *Tol2* family elements have undergone at least five horizontal transfer events. This number represents the minimum estimate. If one or more of these transfer events involved intermediate host species, the actual number of horizontal transfer events would be greater.

### 5. Model for investigating the survival of DNA-based transposable elements

Until now, *Tol2* and related elements have served as an excellent model for studying the evolution of DNA-based transposable elements in teleost fishes because autonomous copies occur naturally in medaka and horizontal transfer between host organisms has been demonstrated (Koga 2000; Ishiyama 2017). In the present study, we identified the *Tel2* element in lampreys and examined both its past chromosomal transposition within the host genome and its horizontal transfer between teleost fishes and lampreys. These findings suggest that *Tol2* family elements may also have been transferred among other vertebrate lineages across the evolutionary boundary between jawless and jawed vertebrates, extending the applicability of this model to encompass all vertebrates.

Our primary approach was a genome database survey. Each database represents the genome of only a single individual. Although we also performed PCR analysis, only four individuals were available for examination. Therefore, it remains possible that an autonomous *Tel2* copy exists in wild lampreys that have not yet been analyzed. Regardless of whether such copies are found, *Tol2* and related elements now provide a unique vertebrate model for testing the widely accepted view of the long-term survival of DNA-based transposable elements through horizontal transfer. As genome databases continue to improve and expand, additional organisms harboring *Tol2* family elements may be identified, leading to a more detailed reconstruction of the stepwise history of their horizontal transfer.

## Materials and Methods

### 1. Online survey of *Tol2*-like sequences

The Basic Local Alignment Search Tool (BLAST) survey was performed on the National Center for Biotechnology Information (NCBI) website (https://blast.ncbi.nlm.nih.gov/Blast.cgi) (Camacho 2009) in December 2025. The query sequences were nucleotides 1–4,682 (full-length), 1–150, or 4,533–4,682 of the medaka *Tol2* element (GenBank: D84375.2). The search parameters were as follows: Database = Core Nucleotide database (core_nt); Organism = lampreys (taxid:7745), *L. fluviatilis* (taxid:7748), *L. planeri* (taxid:7750), or not specified; Optimize for = Highly similar sequences (megablast). All hits were regarded as sequences similar to the queries. The hit sequences were mapped onto the chromosomes of the *L. fluviatilis* and *L. planeri* genome assemblies (PRJEB77186 and PRJEB77190, respectively). All candidate sequences identified in this survey are listed in Tables S2 and S3. The presence of the *Tol2*-like sequences in the alternative haplotype assemblies was confirmed using datasets PRJEB77117 (*L. fluviatilis*) and PRJEB77191 (*L. planeri*).

### 2. Searching open reading frames (ORFs)

The ORF search was performed using the ORF finder tool (https://www.ncbi.nlm.nih.gov/orffinder/). The complete *Tel2* sequences, spanning from the left TIR to the right TIR, were used as query sequences. The search parameters were as follows: Minimum ORF length = 300 nt; Genetic code = 1 (Standard); ORF start codon = Any sense codon.

### 3. Comparison of sequences

Sequence alignment was performed using the multiple sequence alignment tool CLUSTALW (https://www.genome.jp/tools-bin/clustalw) (Thompson 1994). The sequences analyzed, consisting of partial coding sequences of the transposase genes or tyrosinase (TYR) genes, were obtained from the NCBI GenBank database (https://www.ncbi.nlm.nih.gov/genbank/) and are listed in Table S6.

### 4. Pairwise *dN*/*dS* analysis

The *dN*/*dS* ratio was calculated using MEGA X (version 10.1.8) software (https://www.megasoftware.net/) (Kumar 2018). The sequences analyzed were the complete coding sequences of the *TYR* gene from each species, obtained from GenBank. Their accession numbers are listed in Table S6.

### 5. Animal experiments

This study was approved by the Ethics Review Board for Animal Experiments at Nagoya University, the Xunta de Galicia, and the University of Vigo Committee for Animal Use in the Laboratory. All animal experiments were conducted in accordance with the relevant regulations, including the Act on Welfare and Management of Animals in Japan (Act No. 105 of October 1, 1973), the Nagoya University Regulations on Animal Care and Use in Research (Regulation No. 74 of April 1, 2020), Directive 2010/63/EU of the European Parliament, and the Spanish regulations on the protection of animals used for scientific purposes (Royal Decree 53/2013).

### 6. Genomic DNA extraction

The *L. fluviatilis* specimens obtained from Sweden were maintained at the University of Vigo, Spain. The animals were anesthetized with 0.01% tricaine methanesulfonate dissolved in water before dissection. Tail tissue fragments were fixed in ethanol and sent to Nagoya University, Japan, for molecular analyses. Medaka (*O. latipes*; himedaka strain) specimens were purchased from Meito Suien (Nagakute, Japan). Tail fin tissues were collected under anesthesia with 0.01% tricaine. Genomic DNA was extracted from each tissue using the DNeasy Blood & Tissue Kit (Cat. No. 69504, QIAGEN). The DNA samples were stored at 4 °C until use.

### 7. PCR

We designed eight primers to amplify *Tol2*-related sequences from *L. fluviatilis* genomic DNA. Their sequences and nucleotide positions are listed in Table S7. All primers were designed based on the medaka *Tol2* sequence and correspond to the Lf-1 copy of the lamprey *Tel2* element. The predicted amplicon lengths for each primer pair are shown in Figure 6A. PCR amplification was performed in a total reaction volume of 10 μL containing genomic DNA at 2 ng/μL, 0.2 μM of each primer, and the remaining reagents as recommended by the manufacturer of KOD-FX Neo (KFX-201, TOYOBO). For primer pairs including P1a or P4b, the PCR conditions were as follows: initial denaturation at 95 °C for 2 min; 40 cycles of 96 °C for 20 s, 50 °C for 15 s, and 72 °C for 30 s; followed by a final extension at 72 °C for 4 min. For the remaining primer pairs, the PCR conditions were as follows: initial denaturation at 94 °C for 2 min; 35 cycles of 98 °C for 10 s, 60 °C for 15 s, and 68 °C for 20 s; followed by a final extension at 68 °C for 60 s. After amplification, 5 μL of each PCR product was mixed with 1 μL of loading dye and subjected to electrophoresis on a 0.8% agarose gel for products amplified with the P1a–P4b primer pair or on a 1.4% agarose gel for products amplified with the other primer pairs.

## Acknowledgments

Dr. Naoyuki Yamamoto helped with the arrangement of the *L. fluviatilis* specimens for PCR analysis. This work was performed with a laboratory budget from Nagoya University.

## Competing interests

The authors declare no competing interest.

## Supplemental figure

**Figure S1.**
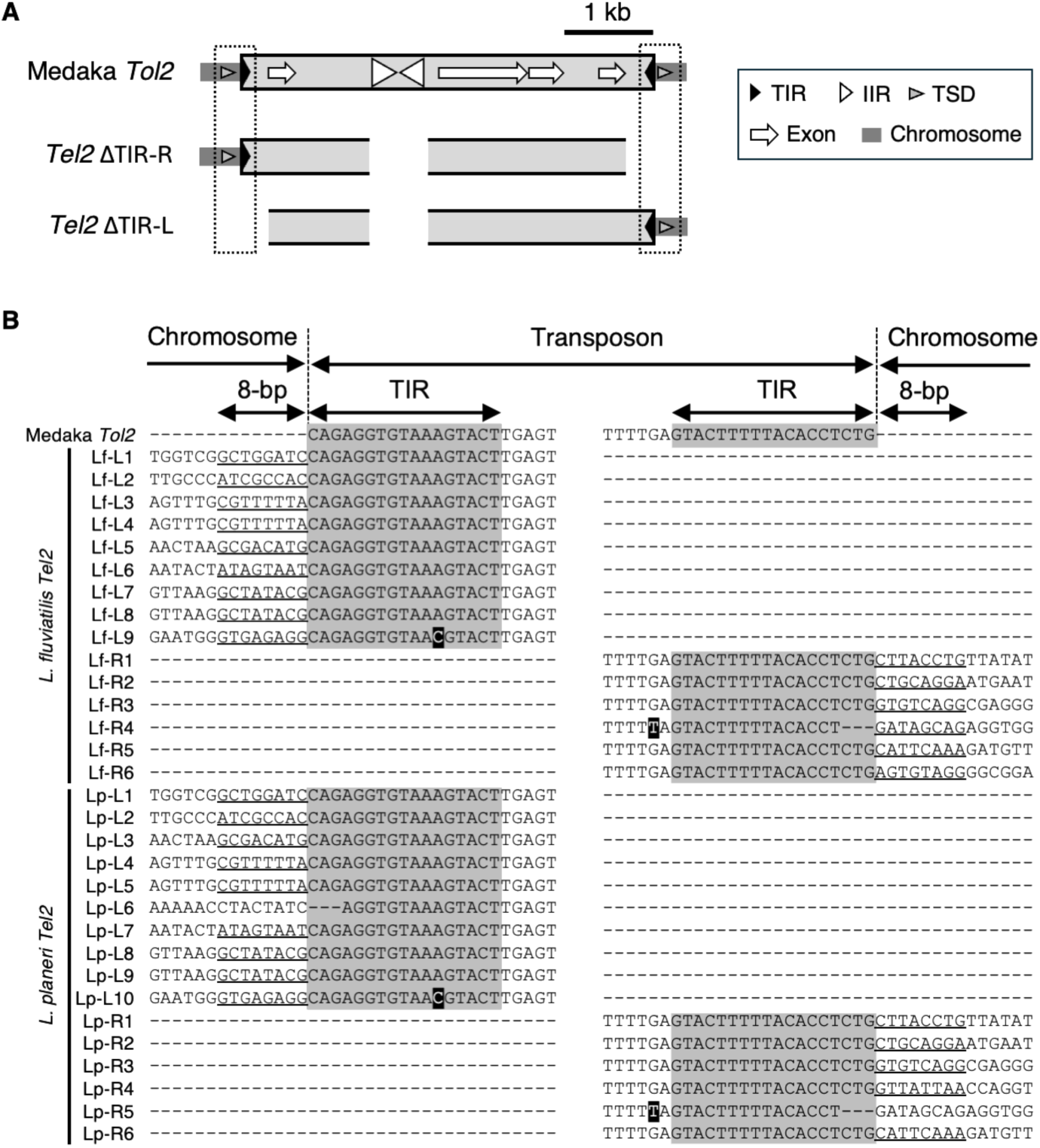
Structures of incomplete *Tel2* termini and adjacent host genomic sequences. **A**. Overview of the incomplete *Tel2* termini and the adjacent host genomic sequences corresponding to the TIRs and TSDs of the *Tol2* element. **B**. Comparison of the terminal sequences of the *Tol2* element and *Tel2* copies containing only one terminal region. Shaded bases indicate nucleotide substitutions relative to *Tol2*. The eight nucleotides immediately adjacent to each *Tel2* TIR are presumed to represent the TSD; however, they were not evaluated because the opposite terminus is absent owing to deletion. This figure provides supplementary information for Figure 3.

## Notes

### Competing Interest Statement

The authors have declared no competing interest.

### Summary of Updates

Revised the whole contents for submission

